# Digital spatial profiling of coronary plaques from persons living with HIV reveals high levels of STING and CD163 in macrophage enriched regions

**DOI:** 10.1101/2020.07.28.221325

**Authors:** Celestine N. Wanjalla, Liang Guo, Daniela T. Fuller, Mona Mashayekhi, Samuel Bailin, Curtis L. Gabriel, Tecla Temu, Jingjing Gong, Yan Liang, Renu Virmani, Aloke V. Finn, Spyros A. Kalams, Simon A. Mallal, Jonathan J. Miner, Joshua A. Beckman, John R. Koethe

**Author notes:** Corresponding authors: Celestine N. Wanjalla, MD, PhD; Division of Infectious Diseases, Vanderbilt University Medical Center, A-2200 MCN, 1161 21st Ave S., Nashville, TN, 37232-2582. (o), (f).

## Abstract

**Background:** Chronic innate and adaptive immune activation may contribute to high prevalence of cardiovascular disease in persons living with HIV (PLWH).

**Methods:** We assessed coronary plaques from deceased PLWH (n=6) and HIV-negative (n=6) persons matched by age and gender. Formalin-fixed, paraffin-embedded 5μm thick sections were processed using Movat, hematoxylin and eosin, immunohistochemical and immunofluorescence stains. Immune cell populations were measured using surface antibodies, and immune-related protein expression from macrophage rich, T-cell rich and perivascular adipose tissue regions using GeoMx^®^ digital spatial profiling.

**Results:** Coronary plaques from PLWH and HIV-negative persons had similar plaque area and percent stenosis. Percent CD163^+^ cells as measured by immunohistochemical staining was significantly higher in PLWH, median 0.29% (IQR 0.11-0.90) vs. 0.01% (IQR 0.0013-0.11) in HIV-negative plaque, p = 0.02 (Figure 1A). Other surface markers of innate cells (CD68 ^+^, p=0.18), adaptive immune cells (CD3^+^, p=0.39; CD4^+^, p=0.09; CD8^+^, p=0.18) and immune trafficking markers (CX3CR1^+^, p=0.09) within the coronary plaque trended higher in HIV-positive plaques but did not reach statistical significance. GeoMx^®^ digital spatial profiling showed higher differential protein expression of CD163 (scavenger receptor for hemoglobin-haptoglobin complex), stimulator of interferon gamma (STING, a cytosolic DNA sensor), CD25 and granzyme-B in the HIV-positive compared to HIV-negative, p<0.05(Figure 1B).

**Conclusions:** Increased inflammation within the coronary plaques of PLWH is characterized by more innate and adaptive immune cells. Higher STING expression in PLWH suggests that immune response to viral antigens within the plaque might be a driver above other stimulants. STING inhibitors are available and could be investigated as a future therapeutic target in PWH if these results are replicated with a larger number of plaques.

**Graphical Abstract:** 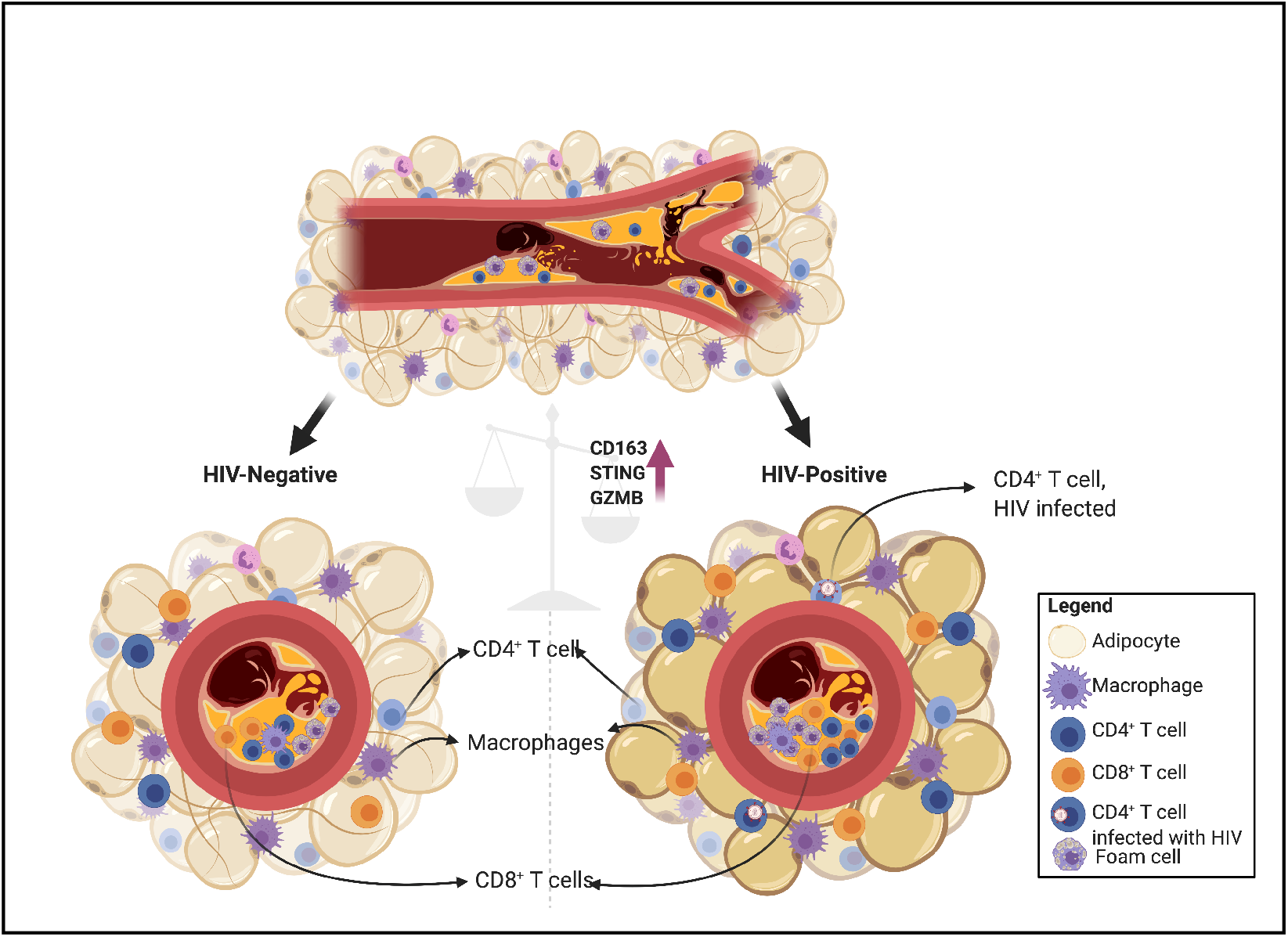

**Highlights:**

1. Immunohistochemical and fluorescent stains combined with GeoMx^®^ digital spatial profiling allowed for deep characterization of immune cells within intact coronary plaques and perivascular adipose tissue
2. Coronary plaques from HIV-positive persons had higher proportion of CD163^+^ immune cells compared to HIV-negative persons
3. Differential protein expression of immune-rich regions of interest within intact 5μm sections of coronary plaques revealed higher levels of stimulator of interferon gamma (STING) in HIV-positive persons

## Introduction

Persons living with human immune deficiency virus (PLWH) have a greater prevalence of cardiovascular disease (CVD) compared to HIV-negative persons, which is not explained by differences in traditional risk factors and persists despite suppression of plasma viremia on antiretroviral therapy (ART).^1–5^ Chronic inflammation due to HIV-infection and other viral pathogens such as cytomegalovirus (CMV) and hepatitis C virus, have been linked to accelerated atherosclerosis^6, 7^ and may in part explain this increased risk. Aortic and carotid artery inflammation measured by 18F-fluorodeoxyglucose positron emission tomography (FDG-PET) imaging,^14, 15^ showed greater tracer uptake in PLWH and was associated with higher plasma hsCRP, IL-6, CX3CR1^+^ monocytes, and potentially CX3CR1^+^ CD4^+^ T cells. ^14^ In general, PLWH on ART have higher levels of inflammation compared to HIV-negative persons.^8, 9^ Higher levels of circulating interleukin-6 (IL-6), high sensitivity c-reactive protein (hsCRP) and d-dimer are associated with CVD events in PLWH^10–12^. Studies in ‘elite controllers’, or PLWH with low or undetectable plasma viremia in the absence of ART, notably also showed higher carotid intimal media thickness compared with HIV-negative controls after adjusting for traditional risk factors, indicating that antiretroviral agents or higher HIV replication per se was not the driver of vascular disease.^13^

Different circulating immune cell subsets have been associated with CVD in the general population and PLWH, though results have been conflicting. CD27^−^ CD28^−^ CD45RO^+^ CD4^+^ T cells, for example, were associated with increased mortality from coronary heart disease.^16^ In contrast, a large analysis combining participants in the Multi-Ethnic Study of Atherosclerosis (MESA) and the Cardiovascular Health Study (CHS) found that circulating lymphocytes (CD4^+^, CD8^+^, CD19^+^) and monocytes (CD14^+^) were not associated with future myocardial infarction in otherwise healthy adults^17^. Among PLWH, lower CD4^+^ T cell counts have been linked with higher rates of non-AIDS diseases, including CVD.^18^ CD4^+^ T cell counts less than 200 cells/mm^3^ were also associated with greater arterial stiffness^19^ and carotid plaque (IMT > 1.5mm).^20^ In other studies and in contrast, higher absolute CD4^+^ T cell counts post-ART at the time of CVD assessment have also been associated with cardiovascular aging in PLWH.^21^ A potential explanation of this paradox is the expansion of a subset of cytotoxic CMV-specific CD4^+^ T cells in the presence of ART which are highly inflammatory and atherogenic. The majority of PLWH are co-infected with CMV and have inflated CD4^+^ and CD8^+^ T cells dedicated to controlling CMV replication. ^22, 23^ A higher percentage of CMV-specific CD8^+^ T cells in PLWH is associated with increased carotid intima-media thickness,^24^ while higher anti-CMV IgG titers are associated with subclinical carotid artery disease and increased mortality from coronary heart disease.^16, 25^

Although inflammation is thought to play an important role in the development of an atheroma, and therapeutic targets including antiviral therapies or immune therapies have been proposed, to our knowledge there has been no research on the immune landscape of arterial plaques in persons with HIV. A comparison of plaques from PLWH vs. HIV-negative persons may delineate inflammatory pathways contributing to a higher burden of CVD. We previously described the C-G-C^+^ CD4^+^ T cells co-expressing CX3CR1 and CD57 in HIV-positive diabetics, and found that these cells are predominantly T_EMRA_ cells and overlap with CD28^−^ CD4^+^ T cells that have been described in aging individuals^26^. Further, a recent publication using single cells isolated from carotid plaque samples of HIV-negative individuals, showed expression of CX3CL1 in nascent plaque and CX3CR1^+^ CD57^+^ CD4^+^ T cells within the plaque by flow cytometry.^27^ They did not include plaque from HIV-positive samples which we address in this study.

Understanding the role of inflammation in atherosclerosis requires definitive evidence using in-depth analysis of immune cells within plaque tissue^28^. This has not been done in PLWH and until now, we were limited in our ability to investigate immune subsets within vulnerable human plaque tissue due to high levels of necrosis that affected the integrity of surrounding tissue and immune cell yield^29–33^. In this paper, we investigated immune cells within coronary plaque tissue and surrounding adventitia using GeoMX^®^ digital spatial profiling which allowed us to pick regions within the plaque and compare surface and intracellular protein expression. Plaques are heterogenous and understanding the immune drivers of atherogenesis is enhanced by the new technologies that allow us to select regions within the plaque for detailed analysis.

As metabolic disease and CVD are co-travelers, we hypothesized that C-G-C CD4^+^ T cells which express CX3CR1 could traffic to inflamed endothelium and contribute to CVD progression in PLWH. C-G-C CD4^+^ T are cytotoxic and more commonly anti-viral. We obtained coronary plaques from twelve deceased individuals with (n=6) and without HIV (n=6) of similar age and sex distribution. The coronary plaques were staged and evaluated for immune cells using immunohistochemistry staining. We found that plaques from PLWH had higher proportions of immune cells per area, with significantly more CD163^+^ cells. Most importantly, using GeoMX^®^ digital spatial profiling, we found a significantly higher expression of stimulator of interferon gamma (STING), CD163, and several immune proteins consistent with a cytotoxic response in the HIV-positive coronary plaque.

## Methods and materials

### Human samples

This study used deidentified human coronary plaque samples/ autopsy specimens approved for exempt review by the institutional review boards of CVPath and Vanderbilt University Medical Center (IRB# 200148). Slides containing major epicardial coronary arteries sectioned at 3-4mm intervals from six HIV-positive and six HIV-negative persons who died of sudden cardiac death were obtained from the CVPath Institute (Gaithersburg, MD) (**Supplementary Table 1**). CVPath Institute maintains curated biorepository of coronary artery beds from over 7000 autopsy hearts from the Office of the Chief Medical Examiner of the State of Maryland (OCME-MD) collected between 2005 and 2019, for providing cardiac consultation. Each heart is evaluated by a cardiac pathologist for staging and the cause of death, if known, is recorded along with de-identified demographic information including age, gender, and race. Each specimen is fixed in 10% formalin and regions of interest are decalcified before processing. The arteries with coronary plaques are fixed and serial sections embedded in paraffin. Sections are cut at 5-6μm and mounted on charged slides^35^.

### Immunohistochemical staining

FFPE sections were stained using Movat pentachrome and hematoxylin and eosin stains (H&E). We selected immune markers to define innate and adaptive Immune cells present in the plaques. These were identified by immunohistochemical staining (IHC) with antibodies against T cells: CD3 (Roche Cat # 790-4341, pre-diluted), CD4 (Roche, Cat# 790-4423, pre-diluted), CD8; macrophages: CD68 (Roche Cat# 790-2931, pre-diluted), CD163 (Leica Cat# NCL-L-CD163, antibody 1:50), vascular cell adhesion molecule 1 (VCAM-1) (Abcam ab134047, 1:500, diluted) and an endothelial homing chemokine receptor, CX3CR1 (Abcam ab8021, 1:1000 dilution), DISCOVERY OmniMap anti-Ms HRP cat # 760-4310 or anti-Rb HRP cat # 760-4311 and developed by the NovaRed kit (Vector Laboratories). The images were captured by Axio Scan.Z1 (Zeiss, Germany) using a 20X objective. IHC staining was quantified in segments with the most severe stenosis using the area quantification module on the HALO image analysis platform (Indica Labs, Corrales, NM) as previously published.^35, 36^

### Digital Spatial profiling of protein expression

Expression of multiple immune-related proteins was measured on 5μm thickness formalin-fixed paraffin-embedded (FFPE) tissue sections from the coronary plaques of two male HIV-positive and HIV-negative with the highest degree of immune cell infiltration. FFPE sections were treated with citrate buffer (pH6) for antigen retrieval. They were bathed in a multiplexed cocktail of primary antibodies with photocleavable DNA-indexing oligos (GeoMX^®^ Immune profile core, Immune Cell typing, Immune Activation Status, IO Drug Target and Pan Tumor modules), fluorescent anti-CD3 (magenta, Cat.# UM500048), anti-CD8 (green, Cat.# 14-0008-82), anti-CD68 (yellow, Cat.# sc-20060) and SYTO 83 nuclear staining (Cat.# S11364). Fluorescence microscopy on the GeoMX^®^ platform was used to image the slides. Twelve regions of interest/areas of interest (ROIs/AOIs) from each were processed using the NanoString’s GeoMX^®^ digital spatial profiling platform (https://www.nanostring.com/scientific-content/technology-overview/digital-spatial-profiling-technology). Images of stained sections were captured at 20X magnification. ROIs/AOIs for molecular profiling were selected as geometric shapes in macrophage-abundant, T-cell abundant and adipose tissue sections. Per protocol, protein staining was repeated twice to verify the results. AOIs were exposed to ultraviolet light (365nm) to release the indexed oligos/barcodes for collection. Oligos were captured from the AOI by microcapillaries and dispensed into 96-well plates. Following collections from all AOIs, the oligos (hybridized to unique four-color, six-spot optical indexing barcodes) were quantified on the nCounter analysis platform. Data were normalized to area; signal-to-noise ratios (SNR) were calculated using isotype controls. Proteins with SNR less than 2 were not included in differential expression analysis. Data were visualized by unsupervised hierarchical clustering. Differential gene expression was analyzed by unpaired *t-test* with Benjamini Hochberg (BH) correction.

### Statistical analysis

Differential expression of proteins in coronary plaques and correlation plots of protein expression were analyzed using *t-test* on the GeoMx^®^ software platform with Benjamini Hochberg correction for multiple comparisons, where applicable. Statistical differences in immune cells within coronary plaques of HIV-positive and HIV-negative persons were calculated using GraphPad Prism 8 and R v.3.6.1.

## Results

### Demographics of HIV-negative and HIV-positive persons

We obtained coronary plaques from six HIV-positive (median age 50) and six HIV-negative deceased persons (median age 52) (**Supplementary Table 1**). Half of each group was female. There were more Caucasians in the HIV-negative group (5/6) compared to the HIV-positive group (1/6). The coronary plaque lesions consisted of early and late atheroma (**Supplementary Table 1**). The cardiac death categories, as applicable, are also provided for reference in the table.

### Coronary plaque morphology and immune cell constituents in HIV-positive and HIV-negative persons

We first compared the coronary plaque histology between HIV-positive and HIV-negative persons. Movat and H&E stains were used to define the coronary plaque constituents; three representative images are shown (**Figure 1A**). There was no significant difference in plaque area (median 4.74E^6^ μm^2^ [2.78E^6^ – 8.00E^6^] in HIV-negative vs. 7.05 E^6^ μm^2^ [5.59E^6^ – 8.81E^6^] in HIV-positive, *p=0.31*) or percent stenosis (median 66% [54 – 72] in HIV-negative vs. 50% [41 – 62] in HIV-positive, *p=0.13*) (**Figure 1B**). Using IHC, we quantified cells of the innate and adaptive immune system. CD68 is a surface marker expressed on monocytes and macrophages; CD3/CD4/CD8 are markers expressed on T cells and vascular cell adhesion molecule 1 (VCAM-1) and CX3CR1) are markers associated with trafficking to inflamed endothelium (**Figure 2A**). We found no difference in the percentage of CD68^+^ cells between HIV-negative (median 0.38% per μm^2^ [0.13, 1.11]) and HIV-positive (1.58% [0.28, 2.94]) coronary plaques, *p=0.18*. Similarly, the percentage of T cells was not different between HIV-negative and HIV-positive coronary plaque: CD3 (median 0.07% per μm^2^ [0.03, 0.23] vs. 0.2% per μm^2^ [0.11, 0.34], *p=0.39*]); CD4 (median 0.001% per μm^2^ [0.0006, 0.05] vs. 0.1% per μm^2^ [0.007, 0.26], *p=0.09*]) or CD8 T cells (median 0.05% per μm^2^ [0.004, 0.17] vs. 0.17% per μm^2^ [0.08, 0.28], *p=0.18*]). Similar trends were seen in the markers associated with trafficking of cells to inflamed endothelium VCAM-1 (median 0.008% per μm^2^ [0.002, 0.03] vs. 0.05%per μm^2^ [0.009, 0.2], *p=0.24*]) and CX3CR1 (median 0.1% per μm^2^ [0.08, 0.9] vs. 0.8% per μm^2^ [0.6, 1.3], *p=0.09*]). In aggregate, there appeared to be a trend towards more immune cells in coronary plaques from HIV-positive individuals (**Figure 2B**). Similarly, we obtained high magnification images of perivascular adipose tissue adjacent to the coronary plaques that showed the presence of innate and adaptive immune cells (**Supplementary Figure 1**)

**Figure 1.**
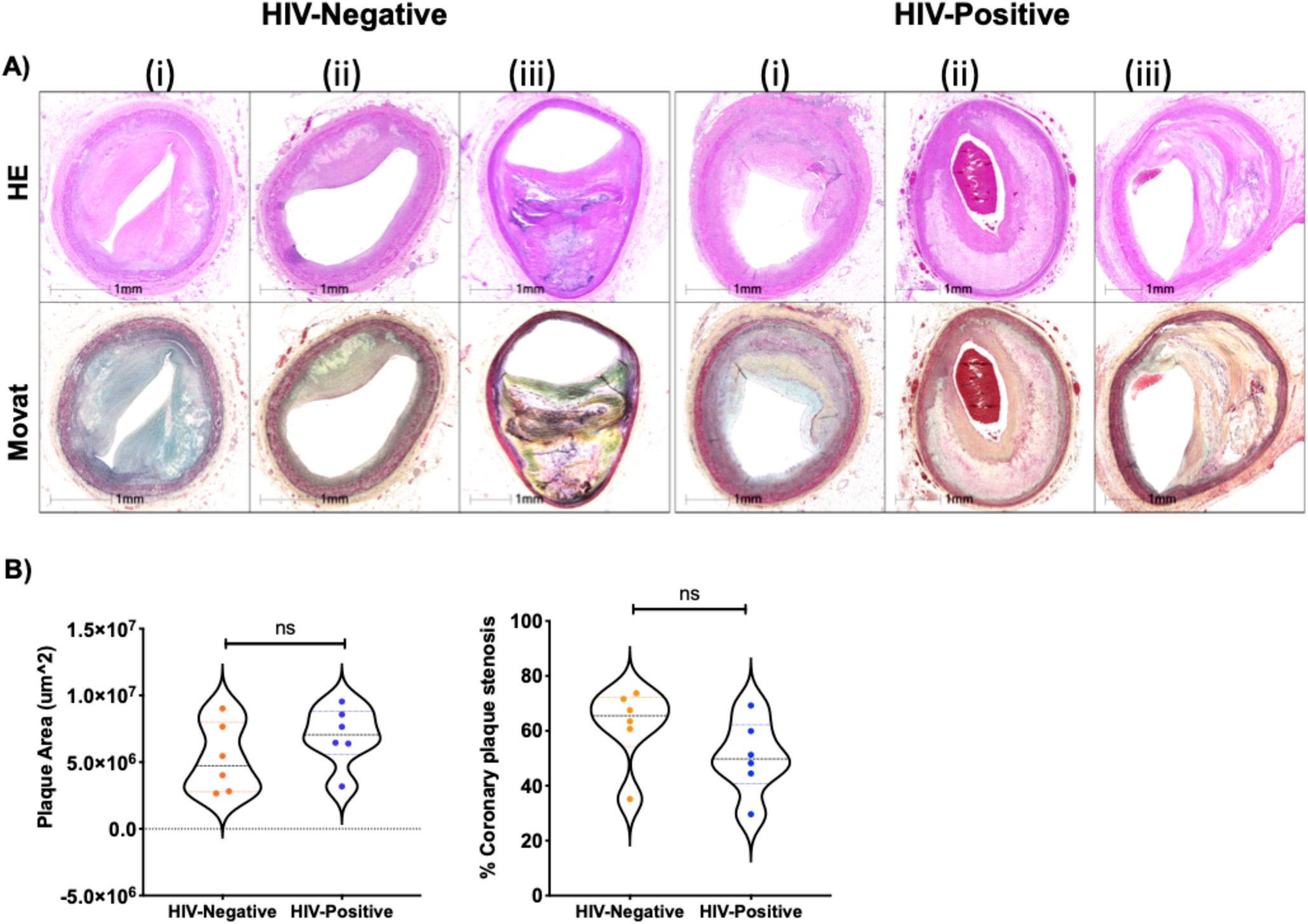
Characterization of coronary plaques. Representative coronary plaques from three HIV-positive and three HIV-negative individuals were stained with H&E and Movat stain (A). Plaque area (μm^2^) and % plaque stenosis were measured and calculated in 6 individuals per group (B). Statistical analysis, Mann-Whitney; *ns not significant*

**Figure 2.**
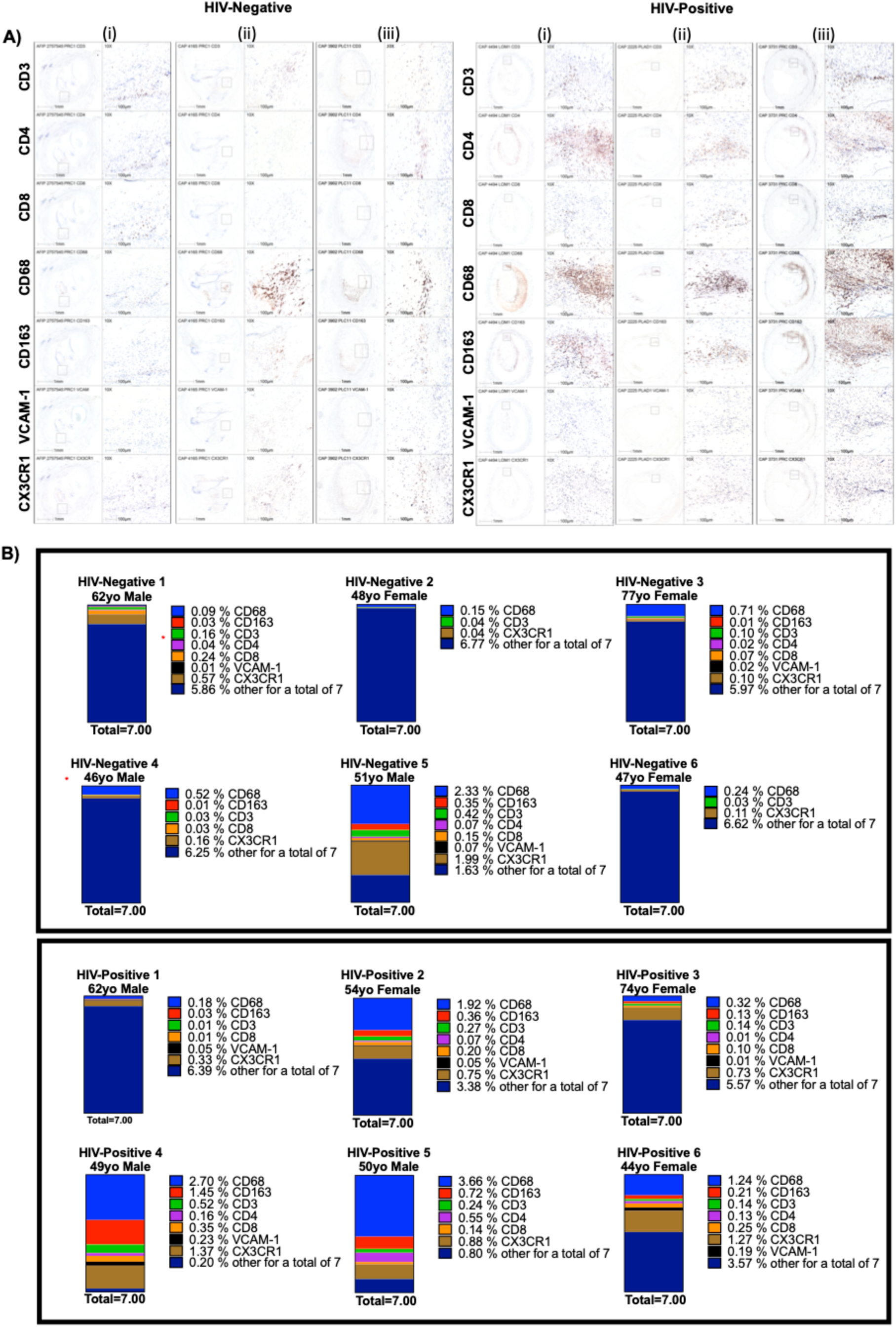
Coronary plaque immune cell constituents in HIV-positive and HIV-negative persons. IHC stains of CD3, CD4, CDS, CD6S, CD163, VCAM-1 and CX3CR1 in three out of six representative coronary plaques from HIV-positive and negative deceased persons (A). The percentage of CD6S^+^, CX3CR1^+^, CDS^+^, CD4^+^, VCAM-1^+^ and CD163^+^ were determined and displayed using a stacked bar chart showing additive percentage of immune cells/um^2^ in each individual. The highest total percentage of cells was ~7%, which was used as the total in in all individuals to allow for direct comparisons (B). Coronary plaques labeled HIV-Negative #5 and HIV-Positive #4 (marked by red*) were selected for digital spatial profiling.

Membrane bound CD163 is a scavenger receptor of hemoglobin-haptoglobin complexes that is expressed exclusively on macrophages^37^. This receptor in humans can be cleaved by the inflammation-inducible enzyme TNF-α converting enzyme (TACE) to generate soluble CD163 that has been shown to be higher in the peripheral blood of PLWH.^38^ Previous studies comparing sCD163 expression found higher levels in HIV+CMV+ compared to HIV+CMV-persons^42^. CD163^+^ macrophages have been associated with a high level of HIF1α expression and plaque progression due to increased plaque angiogenesis and plaque vulnerability. We found that CD163^+^ cells were more prevalent in the HIV-positive (median 0.29% per μm^2^ [0.11, 1.45]) versus HIV-negative (median 0.01%per μm^2^ [0.001, 0.11]), *p = 0.02*) (**Figure 2**). Although vascular adhesion molecule-1 (VCAM-1^+^) was higher in HIV-positive (median 0.05% per μm^2^ [0.009, 0.20]) versus HIV-negative (median 0.008% per μm^2^ [0.002, 0.03]), *p = 0.24*), this difference was not statistically different in this small sample. Taken together, coronary plaque samples from HIV-positive and HIV-negative deceased persons had a median plaque area and stenosis that was similar. However, there was a trend towards higher percentages of immune cells in HIV-positive samples, with significantly higher CD163^+^ cells and a trend towards higher CX3CR1^+^ cells.

### Innate and adaptive immune cells did not correlate with plaque stenosis in HIV-positive samples

Immune cell enrichment in coronary plaques is non-stochastic and is a harbinger of atherosclerosis progression. Monocytes are recruited early after endothelial injury and are stimulated to become macrophages in the sub-endothelium. Using correlation matrices, we looked to see if there were relationships between plaque area, plaque stenosis and plaque resident immune subsets. Spearman’s rank correlation analysis agnostic to HIV-status, showed that CD68^+^ cells were positively correlated with CD163^+^ (rho=0.79, *p<0.01*), CD3^+^ (rho=0.69, *p<0.05*), CD4^+^ (rho=0.74, *p < 0.01*), and CX3CR1^+^ (rho = 0.71, *p < 0.05*) cells. CD8^+^ (rho = 0.52, *p = 0.09*) and VCAM-1^+^ (rho = 0.42, *p = 0.18*) were not significant (**Figure 3A**). We observed that CX3CR1^+^ cells were positively correlated with CD163^+^ (r=0.88, *p < 0.001*), CD3^+^ (r=0.80, *p < 0.01*), CD4^+^ (r=0.76, *p < 0.01*), CD8^+^ (r= 0.80, *p < 0.01*), and VCAM-1 (r=0.67, *p < 0.05*) (**Figure 3A**). When we stratified samples by HIV-status, percent stenosis was positively correlated with CD163^+^, CD3^+^, CD4^+^, and CD8^+^ cells in HIV-negative (**Figure 3B,D**). On the contrary, percent stenosis was not correlated with all immune subsets in HIV-positive individuals (**Figure 3C,D**).

**Figure 3.**
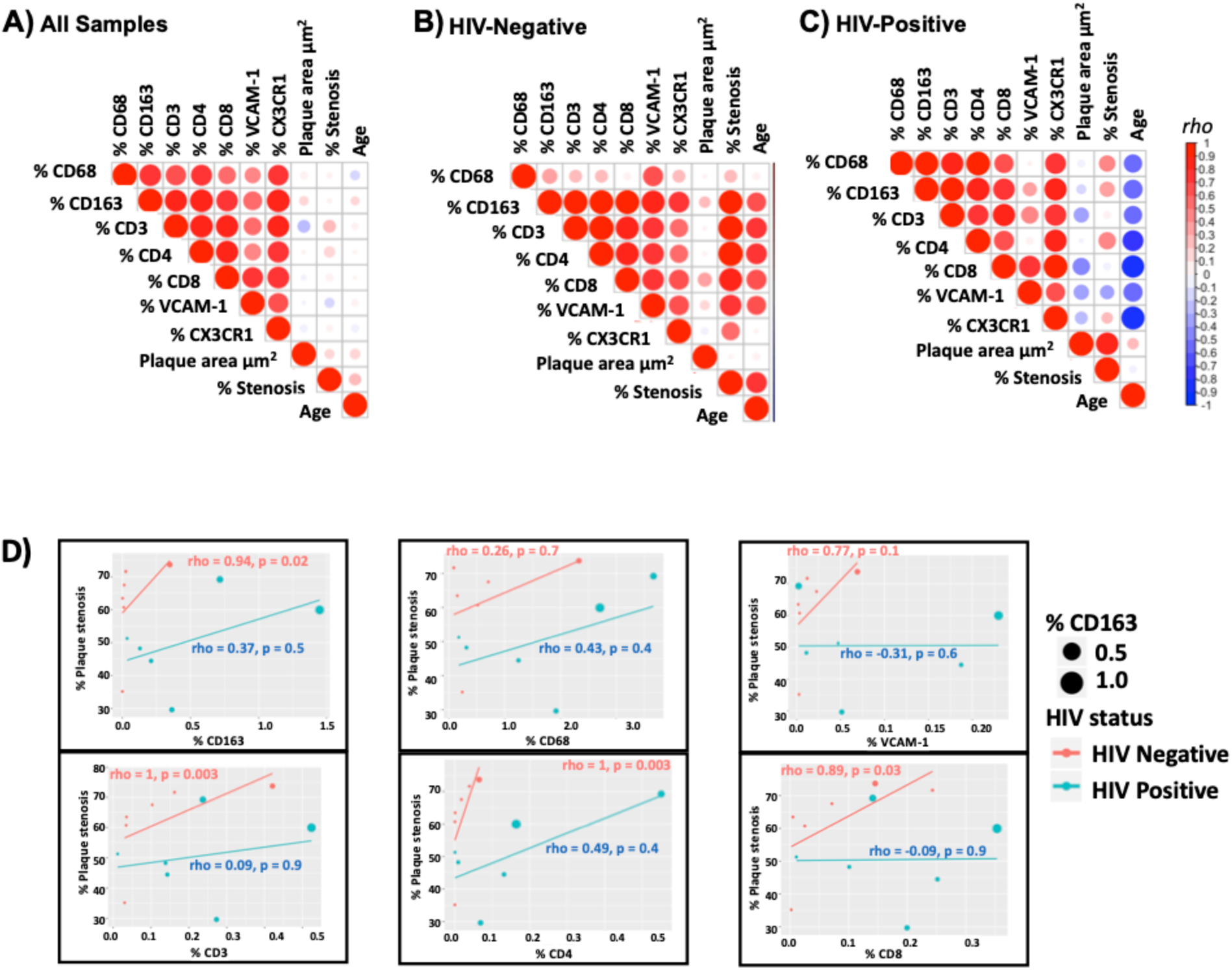
There is a positive correlation between plaque stenosis and CD163, CD3, CD4 and CDS in HIV-negative coronary samples and not HIV-positive. Correlation matrices between immune cells in all samples (A), HIV-positive (B) and HIV-negative (C) were generated in r using the corrplot package. Spearman rank correlation analysis of plaque stenosis and each of the immune subsets was also done on r, the size of the dots represent the percentage of CD163^+^ cells in that specific sample (D). Statistical analysis, spearman correlation analysis, P **<** 0.05 significant.

### Coronary plaque heterogeneity in HIV-positive and HIV-negative persons

Coronary plaque FFPE sections from two individuals (one HIV-positive and one HIV-negative) were selected and matched on age, morphology, sex and percent of immune cells as seen by IHC (CD68^+^ cells, 2.7% HIV-positive and 2.3% in HIV-negative; CX3CR1^+^, 1.4% HIV-positive and 2.0% in HIV-negative). CD163 and VCAM-1 positivity was higher in the HIV-positive samples (IHC stains) compared to the HIV-negative samples (**Supplementary Table 2**). Representative sections from each of these individuals were stained with fluorescently tagged antibodies (CD3, CD8, CD68 and DAPI) and sequential areas of interest (AOI) from each individual was selected (**Figure 4A**). Twelve regions were selected per sample representing regions within the plaque, adventitia, and perivascular adipose tissue (**Figure 4B**). There was significant heterogeneity in areas within the plaque as some regions had predominantly macrophages and others had predominantly CD3^+^/CD8^+^ cells (**Figure 4C**). Images with single fluorescent antibodies were included to show the macrophage and T cells within each AOI (**Supplementary Figure 2**). In general, there was similar degree of heterogeneity in the HIV-positive and HIV-negative plaques.

**Figure 4.**
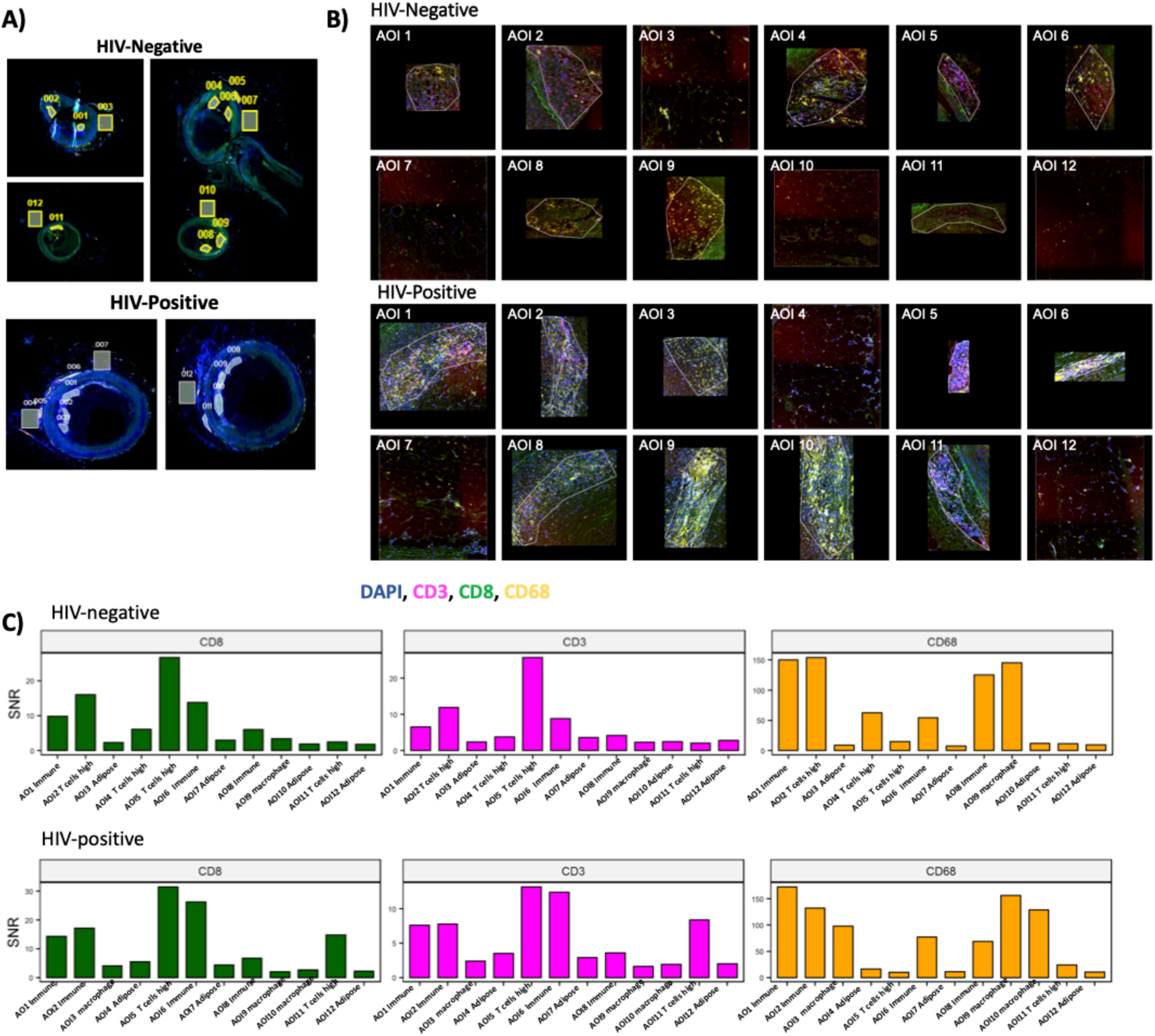
Digital spatial profiling of immune-related proteins in coronary plaques from HIV-positive and HIV-negative individuals. Twelve ROI per sample (HIV-positive and negative) were selected for analysis. Geometric regions of varying sizes were used to select regions of interest within the plaque, in the adventitia and perivascular adipose tissue. (A). Each sample was stained with anti-CD3 (pink), anti-CD8 (green) and anti-CD68 (yellow). SYTO 83 nuclear staining was included to visualize all cells. The twelve different regions selected per sample are designated as area of interest (AOI 1-12) are shown (B). Signal to noise ratio normalized barcode counts of the three fluorescently tagged markers across all 12-AOls demonstrate heterogeneity of coronary plaques in both groups (C).

Using an optical barcode microscope, we obtained digital spatial protein expression data from the coronary plaques of an HIV-positive and HIV-negative person. Molecular profiling using a heatmap showed that adipose tissue AOIs had similar protein expression profiles independent of HIV-status (**Figure 5A**). Differential protein expression of all AOIs by HIV-status showed higher Stimulator of interferon genes (STING), CD163, V-domain immunoglobulin suppressor of T cell activation (VISTA), Bcl-2, Ki-67 and cytotoxic T-lymphocyte-associated protein 4 (CTLA-4) (*p < 0.05*) in the HIV-positive coronary plaque (**Figure 5B-C**).

**Figure 5.**
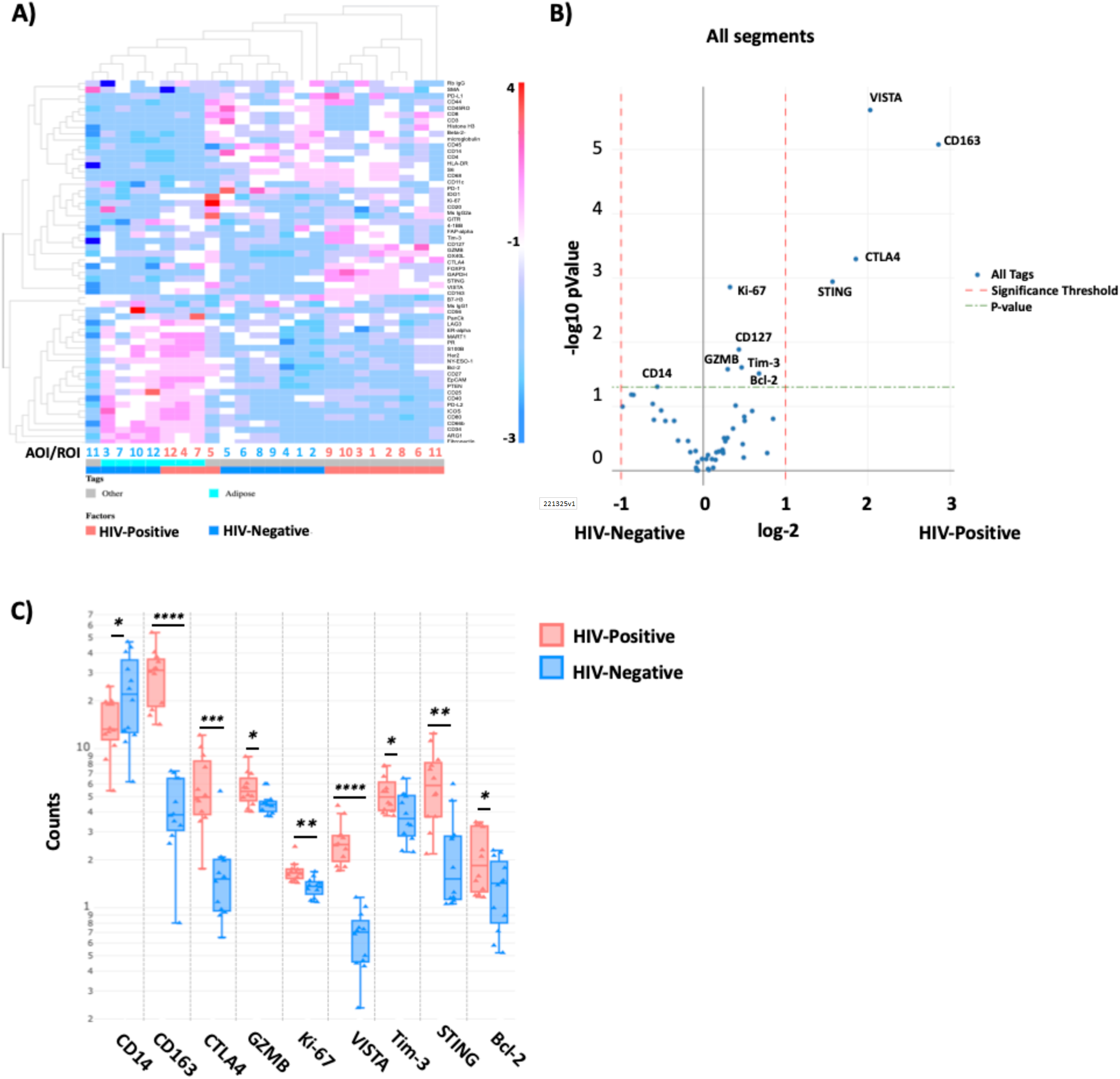
Digital spatial imaging analysis shows clustering of adipose tissue segments independent of HIV-status. A dendrogram grouping area of interest (AOI) segments that have similar protein expression (A). Differential protein expression of all segments by HIV-status (B). Box plots showing STING, VISTA, CD127, Bcl-3, Ki-67 and CD163, higher in HIV-positive and CD14, higher in HIV-negative AOl’s (C). Significance determined by t-test, and BH correction p < *0.05* significant.

### STING is highly expressed in macrophage-rich HIV-positive AOIs

We excluded the adipose tissue and adventitia (external AOIs) and analyzed the differential protein expression of AOIs within the coronary plaque by HIV-status. These included 6 AOIs from the HIV-positive plaque and 7 AOIs from the HIV-negative. Differential protein expression by *t-test* showed significantly higher expression of STING, CD163, VISTA, GZMB, Ki-67, Bcl-2, CD25, Tim-3, CD127 and CTLA4 in the HIV-positive coronary plaque AOIs (**Figure 6A**), while HLA-DR, CD14 was higher in the HIV-negative coronary plaque. The distribution of the different AOIs contributing to these main genes are shown by boxplot (**Figure 6B**). Due to the heterogeneity within the coronary plaques, we used the trend line to show variations in counts by section in both the HIV-positive and negative. CD163 was highly expressed in all AOIs (**Figure 6C**). However, the segments that had the highest proportion of macrophages in the HIV-positive sample (AOI 9 and 10) had the highest expression of STING. Notably although lower expression of STING was present in the HIV-negative plaque, the regions with the highest expression were also those with more macrophages (AOI 1,2 and 6). STING correlated with activation (CD25, R^2^ = 0.77; Tim-3, R^2^ = 0.83), naïve and memory T cells (CD127, R^2^ = 0.68), macrophages (CD163, R^2^ = 0.62). VISTA protein, which is associated with myeloid activation and is a checkpoint inhibitor, expression was correlated with CD163 R^2^ = 0.82 and GZMB R^2^ = 0.71 (**Figure 6C**).

**Figure 6.**
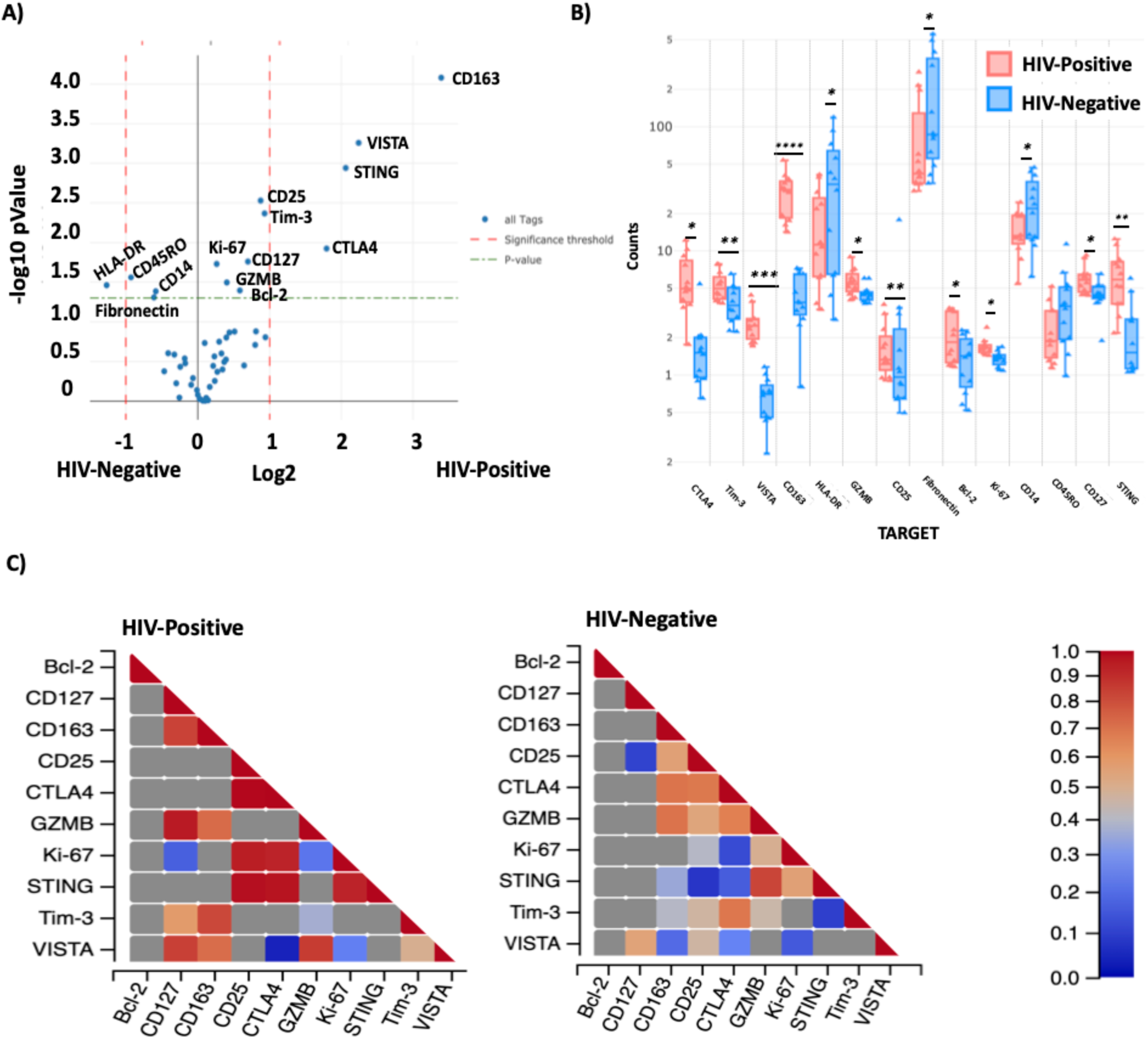
STING, CD163 and VISTA protein expression levels are higher in HIV-positive coronary plaque AOl’s. Violin plot showing differential protein expression between HIV-positive and HIV-Negative AOls within the plaque (A). Box plot showing the median counts of the proteins that were significantly higher in HIV-negative and positive samples (B). There is a positive correlation between STING protein expression and Ki-67 both HIV-positive and HIV-negative coronary plaque samples, and CD25, CTLA4 in the HIV-positive (C). Statistical analysis, t-test with BH correction, (green dashed line) *p* < *0.05* and red dashed lines (fold change > 1).

Analysis of segments external to the plaque (adipose and adventitia) showed higher levels of B7-H3, fibronectin and CD34 expression in HIV-negative AOIs. Heatmap of external segments alone showed similarity in protein levels that clustered based on adipose tissue versus adventitia (**Figure 7A**). Differential protein levels by *t-test* showed higher B7-H3, CD34 and fibronectin in HIV-negative samples (**Figure 7B**). These proteins were highly prevalent in the adipose tissue segments (AOI 3,7,10 and 12) and not the immune T cell rich segment in the adventitia (AOI 5). CTLA4, STING and VISTA were found in high levels in immune cell rich AOIs in the adventitia (AOI 5 and 11) while CD163 was higher in an adipose AOI (4) and adventitia AOI with macrophages and T cells (6) (**Figure 7C).**

**Figure 7.**
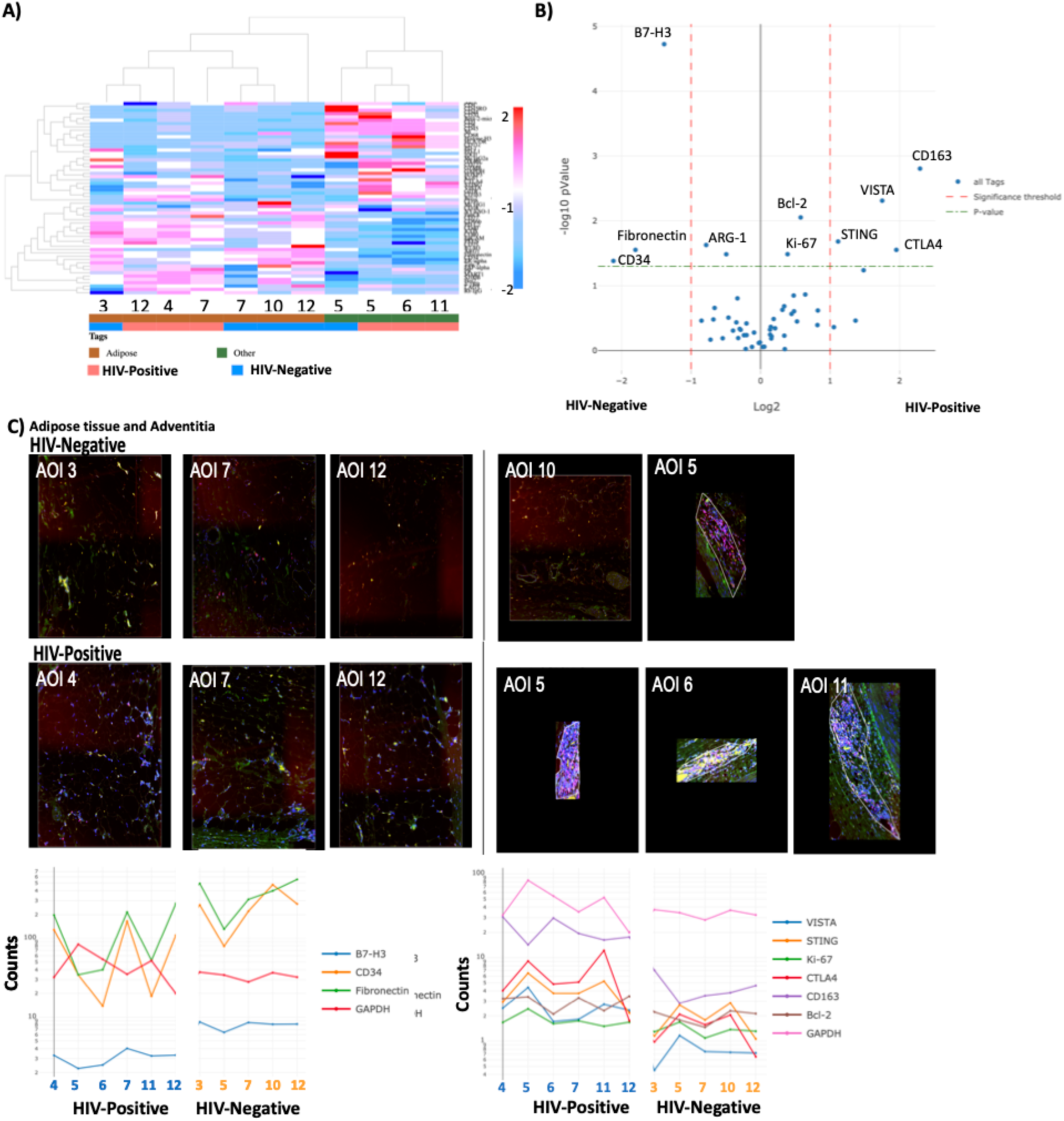
CD163, CTLA4, STING and VISTA are highly expressed in external AOls in the HIV-positive coronary plaque sample. Dendrogram with unsupervised clustering showing protein expression in HIV-positive and HIV-negative AOls (A). Differential gene expression of proteins in external AOls (adipose tissue and adventitia) by HIV-status (B). Trendlines and corresponding external images of the external AOls (C). Statistical analysis, t-test (green dashed line) p < *0.05* and red dashed lines (fold change> 1).

CMV seroprevalence is significantly higher in PWH compared to HIV-negative individuals.^50^ It has been proposed that PWH have a higher level of viral replication within tissue compartments even in the absence of active viremia. Although DNA viruses are the main stimulators of STING via cGAS, RNA viruses such as HIV may also stimulate STING via the retinoic acid-inducible gene I (RIG-I) pathways.^51^ RIG-I is a cytosolic pattern recognition receptor that recognizes double stranded viral RNA. Our proposed hypothesis is that viral DNA and RNA in PLWH, possibly in combination with modified oxidized peptides are transiently activating STING-related pathways within coronary plaques of PLWH, leading to higher levels of inflammation and increased plaque instability (**Figure 8**).

**Figure 8.**
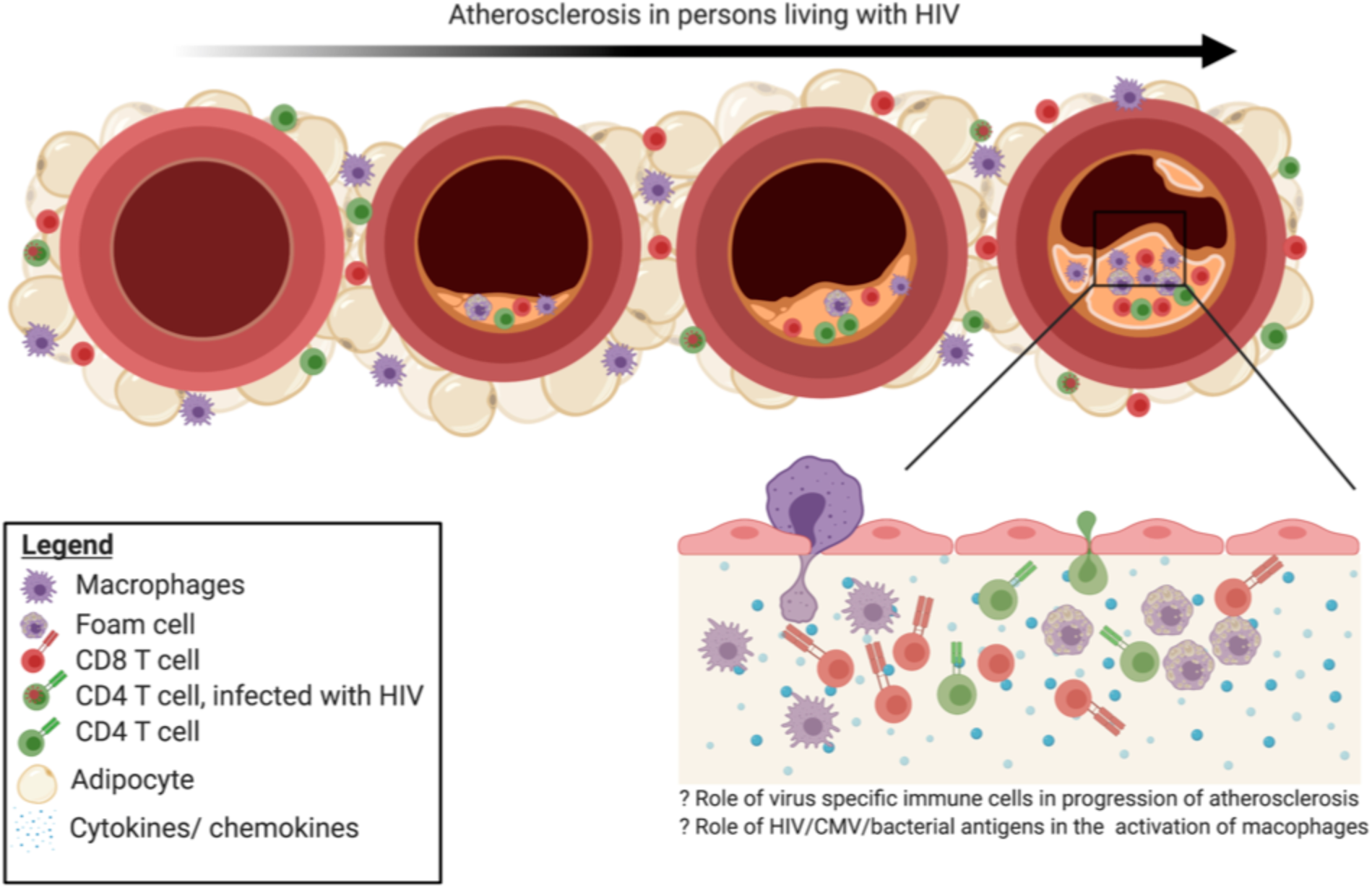
Proposed model. Virus-specific immune responses are important in the progression of atherosclerosis in PLWH. CD4^+^ T cells with HIV DNA have been detected in adipose tissue of PLWH.^28,29,44^ CMV transcripts have also been reported in adipose tissue.^45^ In this paper, we show that PLWH had higher proportions of CX3CR1^+^ CD4^+^ T cells in the perivascular adipose tissue and coronary plaque. A large proportion of CMV-specific CD4^+^ T cells are CX3CR1 ^+^. ^22,36,46,47^ This suggests that virus-specific immune cells are present within early atheromas. Higher expression of STING, CD25, Bcl-2, Ki-67, CD163 and GZMB suggests higher levels of activation in the samples from PLWH compared to the HIV-negative samples. Future studies looking at coronary plaques at different stages of atheroma will be evaluated to answer the question whether low level viral replication (HIV and CMV) can be detected in macrophages/ T cells in coronary plaques of PLWH on ART and whether the magnitude of viral transcripts correlate with the level of immune cell infiltration or activation.

## DISCUSSION

In this study, we hypothesized that coronary plaques from persons with HIV would have a higher proportion of immune cells compared to HIV-negative. Furthermore, they would have an immune profile that is consistent with stimulation by virus compared to coronary plaques from HIV-negative individuals. Newer single-cell analysis has facilitated the investigation of the heterogenous populations of cells present in atherosclerotic plaques.^30^ Understanding the immune components is fundamental if we expect to develop immunotherapies that significantly reduce atherosclerosis progression and CV events. Using GeoMX^®^ digital profiling, we were able to select regions that were enriched with macrophages, T cells, a combination (macrophages and T cells) or perivascular adipose tissue. In general, the coronary plaque AOIs from the representative HIV-positive plaque had higher expression of STING compared to the HIV-negative plaque AOIs. This was even more pronounced in the AOIs that were macrophage rich.

Using coronary samples from HIV-positive and HIV-negative individuals, we quantified the innate and adaptive immune cells within the coronary plaques and perivascular fat. We found a higher frequency of total immune cells within coronary plaques of HIV-positive persons. CD163^+^ cells were significantly higher by IHC and protein expression using GeoMX^®^ digital profiling. Notably, CD163 expression on monocytes and macrophages has been shown to confer susceptibility of infection with both DNA^39^ and RNA^40^ viruses. There are no studies to date that have looked at whether CD163+ macrophages are more permissive to CMV infection. HIV infected CD163^+^ CD68^+^ macrophages have been reported in gut biopsies from untreated PLWH while CMV was primarily detected in epithelial cells^41^. Ongoing studies using RNAscope will define the viral burden - HIV and CMV - within plaque tissue and in perivascular adipose tissue of PLWH to better understand the tissue pathology of these viruses in this context.

STING is an endoplasmic reticulum adaptor protein that can bind DNA viruses and intermediate DNA transcripts of RNA viruses initiating an innate immune inflammatory cascade leading to activate of type I interferons.^52, 53^ The DNA viruses include CMV, Epstein Barr virus (EBV) and herpes simplex virus (HSV). STING is a critical signaling molecule that is involved in tissue inflammation and has been shown to trigger metabolic stress-induced endothelial inflammation in a mouse model. STING agonists have been shown to activate cells that were latently infected with simian immunodeficiency virus (SIV) and enhanced SIV-specific responses *in vivo*^54^. It is possible that a similar effect would be seen with HIV. Thus, in individuals co-infected with HIV and CMV, re-reactivation of CMV might create a setting where HIV infection is enhanced with apoptosis of those CD4^+^ T cells. Notably, CMV-specific CD4^+^ T cells express CX3CR1, and a subset of these CMV-specific cells are protected from HIV-infection^55^. As a result, the CMV-specific cells are likely to accumulate with time in the setting of low levels of CMV replication in tissues. We have previously shown that CMV-specific CD4^+^ T cells were largely T effector memory cells RA^+^ revertant (T_EMRA_) and T effector memory cells (T_EM_); cytotoxic with notable expression of GZMB and perforin at baseline.^22^ To our knowledge, the role of STING in atherosclerosis has not been reported by other groups. Our next steps are to perform a similar analysis on a larger number of samples and define the pathogen burden (both viral and bacterial) within coronary plaques of PLWH and HIV-negative. This will involve *in situ* staining as well as droplet digital PCR to quantify DNA extracted from these samples. Identification of replicating virus in the plaque can provide evidence for the need of CMV-specific antivirals in this patient population or targeted inhibitors of the STING pathway.

This study has several limitations. The deceased persons were confirmed HIV-positive and HIV-negative. However, we do not have information on their ART regimens or viral load at the time of death. Therefore, if we find virus in their coronary plaques, we will not be able to relate the viral burden to their ART compliance. Furthermore, the samples are processed after examination by the medical examiner, therefore there could be a slight delay from when the patient died. Future studies using samples from living donors that are processed within 30 minutes to 1 hour of obtaining the sample are underway. Finally, for the digital spatial profiling, we chose the individuals with the highest proportion of cells so that they would be comparable. Some sections of the HIV-negative blood vessel had the appearance of a total chronic occlusion. Based on our pathology analysis, we opted to match them based on the abundance of immune cells. Future studies will include a larger number of participants and different stages of atherosclerosis.

## Author contributions

Conceptualization, C.N.W., L.G., J.R.K., J.B., S.A.M.; Methodology, C.N.W., L.G., D.T.F., L.M., M.J.T., R.V., Y.L., J.G., A.V.F., S.B., C.L.G., M.M., S.A.M., J.B., J.R.K.; Statistics, C.N.W., J.G., L.M., J.R.K.; Formal Analysis, C.N.W., L.G., D.T.F., L.M., S.B., M.J.T., J.R.K.; Investigation, C.N.W., C.M.W., J.R. K; Resources, J.R.K., J.B., M.J.T., A.V.F., R.V., S.A.K., S.A.M.; Data Curation, C.N.W., L.G., D.T.F., L.M., M.J.T., J. R. K.; Writing – Original Draft, C.N.W., L.G., J.R.K.; Writing – Review & Editing, all authors; Visualization, C.N.W., L.G., D.T.M., M.J.T., J.R.K.; Supervision, J.R.K., L.G., R.V., A.V.F., M.J.T., S.A.M., S.A.K.; Project Administration, J.R.K., J. B., S.A.K.; Funding Acquisition, C.N.W., A.V.F., M.J.T., J.R.K.

## Bibliography

1. Shah ASV, Stelzle D, Lee KK, Beck EJ, Alam S, Clifford S, Longenecker CT, Strachan F, Bagchi S, Whiteley W, Rajagopalan S, Kottilil S, Nair H, Newby DE, McAllister DA, Mills NL. Global burden of atherosclerotic cardiovascular disease in people living with hiv. Circulation. 2018;138:1100–1112

2. Currier JS, Taylor A, Boyd F, Dezii CM, Kawabata H, Burtcel B, Maa JF, Hodder S. Coronary heart disease in hiv-infected individuals. J Acquir Immune Defic Syndr. 2003;33:506–512

3. Triant VA, Lee H, Hadigan C, Grinspoon SK. Increased acute myocardial infarction rates and cardiovascular risk factors among patients with human immunodeficiency virus disease. J Clin Endocrinol Metab. 2007;92:2506–2512

4. Hsue PY, Deeks SG, Hunt PW. Immunologic basis of cardiovascular disease in hiv-infected adults. J Infect Dis. 2012;205 Suppl 3:S375–382

5. Freiberg MS, Chang CC, Kuller LH, Skanderson M, Lowy E, Kraemer KL, Butt AA, Bidwell Goetz M, Leaf D, Oursler KA, Rimland D, Rodriguez Barradas M, Brown S, Gibert C, McGinnis K, Crothers K, Sico J, Crane H, Warner A, Gottlieb S, Gottdiener J, Tracy RP, Budoff M, Watson C, Armah KA, Doebler D, Bryant K, Justice AC. Hiv infection and the risk of acute myocardial infarction. JAMA Intern Med. 2013;173:614–622

6. El-Sadr WM, Lundgren J, Neaton JD, Gordin F, Abrams D, Arduino RC, Babiker A, Burman W, Clumeck N, Cohen CJ, Cohn D, Cooper D, Darbyshire J, Emery S, Fätkenheuer G, Gazzard B, Grund B, Hoy J, Klingman K, Losso M, Markowitz N, Neuhaus J, Phillips A, Rappoport C, Group SfMoATSS. Cd4+ count-guided interruption of antiretroviral treatment. N Engl J Med. 2006;355:2283–2296

7. Pothineni NVK, Subramany S, Kuriakose K, Shirazi LF, Romeo F, Shah PK, Mehta JL. Infections, atherosclerosis, and coronary heart disease. Eur Heart J. 2017;38:3195–3201

8. Armah KA, McGinnis K, Baker J, Gibert C, Butt AA, Bryant KJ, Goetz M, Tracy R, Oursler KK, Rimland D, Crothers K, Rodriguez-Barradas M, Crystal S, Gordon A, Kraemer K, Brown S, Gerschenson M, Leaf DA, Deeks SG, Rinaldo C, Kuller LH, Justice A, Freiberg M. Hiv status, burden of comorbid disease, and biomarkers of inflammation, altered coagulation, and monocyte activation. Clin Infect Dis. 2012;55:126–136

9. Hunt PW, Brenchley J, Sinclair E, McCune JM, Roland M, Page-Shafer K, Hsue P, Emu B, Krone M, Lampiris H, Douek D, Martin JN, Deeks SG. Relationship between t cell activation and cd4+t cell count in hiv-seropositive individuals with undetectable plasma hiv rna levels in the absence of therapy. J Infect Dis. 2008;197:126–133

10. Duprez DA, Neuhaus J, Kuller LH, Tracy R, Belloso W, De Wit S, Drummond F, Lane HC, Ledergerber B, Lundgren J, Nixon D, Paton NI, Prineas RJ, Neaton JD, Group ISS. Inflammation, coagulation and cardiovascular disease in hiv-infected individuals. PLoS One. 2012;7:e44454

11. De Luca A, de Gaetano Donati K, Colafigli M, Cozzi-Lepri A, De Curtis A, Gori A, Sighinolfi L, Giacometti A, Capobianchi MR, D’Avino A, Iacoviello L, Cauda R, D’Arminio Monforte A. The association of high-sensitivity c-reactive protein and other biomarkers with cardiovascular disease in patients treated for hiv: A nested case-control study. BMC Infect Dis. 2013;13:414

12. Triant VA, Meigs JB, Grinspoon SK. Association of c-reactive protein and hiv infection with acute myocardial infarction. J Acquir Immune Defic Syndr. 2009;51:268–273

13. Hsue PY, Hunt PW, Schnell A, Kalapus SC, Hoh R, Ganz P, Martin JN, Deeks SG. Role of viral replication, antiretroviral therapy, and immunodeficiency in hiv-associated atherosclerosis. AIDS. 2009;23:1059–1067

14. Tawakol A, Ishai A, Li D, Takx RA, Hur S, Kaiser Y, Pampaloni M, Rupert A, Hsu D, Sereti I, Fromentin R, Chomont N, Ganz P, Deeks SG, Hsue PY. Association of arterial and lymph node inflammation with distinct inflammatory pathways in human immunodeficiency virus infection. JAMA Cardiol. 2017;2:163–171

15. Yarasheski KE, Laciny E, Overton ET, Reeds DN, Harrod M, Baldwin S, Dávila-Román VG. 18fdg pet-ct imaging detects arterial inflammation and early atherosclerosis in hiv-infected adults with cardiovascular disease risk factors. J Inflamm (Lond). 2012;9:26

16. Spyridopoulos I, Martin-Ruiz C, Hilkens C, Yadegarfar ME, Isaacs J, Jagger C, Kirkwood T, von Zglinicki T. Cmv seropositivity and t-cell senescence predict increased cardiovascular mortality in octogenarians: Results from the newcastle 85+ study. Aging Cell. 2016;15:389–392

17. Olson NC, Sitlani CM, Doyle MF, Huber SA, Landay AL, Tracy RP, Psaty BM, Delaney JA. Innate and adaptive immune cell subsets as risk factors for coronary heart disease in two population-based cohorts. Atherosclerosis. 2020;300:47–53

18. Baker JV, Peng G, Rapkin J, Abrams DI, Silverberg MJ, MacArthur RD, Cavert WP, Henry WK, Neaton JD, (CPCRA) TBCPfCRoA. Cd4+ count and risk of non-aids diseases following initial treatment for hiv infection. AIDS. 2008;22:841–848

19. Ho JE, Deeks SG, Hecht FM, Xie Y, Schnell A, Martin JN, Ganz P, Hsue PY. Initiation of antiretroviral therapy at higher nadir cd4+ t-cell counts is associated with reduced arterial stiffness in hiv-infected individuals. AIDS. 2010;24:1897–1905

20. Hsue PY, Lo JC, Franklin A, Bolger AF, Martin JN, Deeks SG, Waters DD. Progression of atherosclerosis as assessed by carotid intima-media thickness in patients with hiv infection. Circulation. 2004;109:1603–1608

21. Medina S, Wessman D, Krause D, Stepenosky J, Boswell G, Crum-Cianflone N. Coronary aging in hiv-infected patients. Clin Infect Dis. 2010;51:990–993

22. Abana CO, Pilkinton MA, Gaudieri S, Chopra A, McDonnell WJ, Wanjalla C, Barnett L, Gangula R, Hager C, Jung DK, Engelhardt BG, Jagasia MH, Klenerman P, Phillips EJ, Koelle DM, Kalams SA, Mallal SA. Cytomegalovirus (cmv) epitope-specific cd4(+) t cells are inflated in hiv(+) cmv(+) subjects. J Immunol. 2017;199:3187–3201

23. Hunt PW, Martin JN, Sinclair E, Epling L, Teague J, Jacobson MA, Tracy RP, Corey L, Deeks SG. Valganciclovir reduces t cell activation in hiv-infected individuals with incomplete cd4+ t cell recovery on antiretroviral therapy. J Infect Dis. 2011;203:1474–1483

24. Hsue PY, Hunt PW, Sinclair E, Bredt B, Franklin A, Killian M, Hoh R, Martin JN, McCune JM, Waters DD, Deeks SG. Increased carotid intima-media thickness in hiv patients is associated with increased cytomegalovirus-specific t-cell responses. AIDS. 2006;20:2275–2283

25. Parrinello CM, Sinclair E, Landay AL, Lurain N, Sharrett AR, Gange SJ, Xue X, Hunt PW, Deeks SG, Hodis HN, Kaplan RC. Cytomegalovirus immunoglobulin g antibody is associated with subclinical carotid artery disease among hiv-infected women. J Infect Dis. 2012;205:1788–1796

26. Koch S, Larbi A, Derhovanessian E, Ozcelik D, Naumova E, Pawelec G. Multiparameter flow cytometric analysis of cd4 and cd8 t cell subsets in young and old people. Immun Ageing. 2008;5:6

27. Chen B, Morris SR, Panigrahi S, Michaelson GM, Wyrick JM, Komissarov AA, Potashnikova D, Lebedeva A, Younes SA, Harth K, Kashyap VS, Vasilieva E, Margolis L, Zidar DA, Sieg SF, Shive CL, Funderburg NT, Gianella S, Lederman MM, Freeman ML. Cytomegalovirus coinfection is associated with increased vascular-homing cd57. J Immunol. 2020

28. Libby P, Ridker PM, Hansson GK. Progress and challenges in translating the biology of atherosclerosis. Nature. 2011;473:317–325

29. Cochain C, Vafadarnejad E, Arampatzi P, Pelisek J, Winkels H, Ley K, Wolf D, Saliba AE, Zernecke A. Single-cell rna-seq reveals the transcriptional landscape and heterogeneity of aortic macrophages in murine atherosclerosis. Circ Res. 2018;122:1661–1674

30. Fernandez DM, Rahman AH, Fernandez NF, Chudnovskiy A, Amir ED, Amadori L, Khan NS, Wong CK, Shamailova R, Hill CA, Wang Z, Remark R, Li JR, Pina C, Faries C, Awad AJ, Moss N, Bjorkegren JLM, Kim-Schulze S, Gnjatic S, Ma’ayan A, Mocco J, Faries P, Merad M, Giannarelli C. Single-cell immune landscape of human atherosclerotic plaques. Nat Med. 2019;25:1576–1588

31. Lebedeva A, Vorobyeva D, Vagida M, Ivanova O, Felker E, Fitzgerald W, Danilova N, Gontarenko V, Shpektor A, Vasilieva E, Margolis L. Ex vivo culture of human atherosclerotic plaques: A model to study immune cells in atherogenesis. Atherosclerosis. 2017;267:90–98

32. Zhou X, Stemme S, Hansson GK. Evidence for a local immune response in atherosclerosis. Cd4+t cells infiltrate lesions of apolipoprotein-e-deficient mice. Am J Pathol. 1996;149:359–366

33. Mach F, Sauty A, Iarossi AS, Sukhova GK, Neote K, Libby P, Luster AD. Differential expression of three t lymphocyte-activating cxc chemokines by human atheroma-associated cells. J Clin Invest. 1999;104:1041–1050

34. Wanjalla CN, McDonnell WJ, Barnett L, Simmons JD, Furch BD, Lima MC, Woodward BO, Fan R, Fei Y, Baker PG, Ram R, Pilkinton MA, Mashayekhi M, Brown NJ, Mallal SA, Kalams SA, Koethe JR. Adipose tissue in persons with hiv is enriched for cd4. Front Immunol. 2019;10:408

35. Kramer MC, Rittersma SZ, de Winter RJ, Ladich ER, Fowler DR, Liang YH, Kutys R, Carter-Monroe N, Kolodgie FD, van der Wal AC, Virmani R. Relationship of thrombus healing to underlying plaque morphology in sudden coronary death. J Am Coll Cardiol. 2010;55:122–132

36. Guo L, Akahori H, Harari E, Smith SL, Polavarapu R, Karmali V, Otsuka F, Gannon RL, Braumann RE, Dickinson MH, Gupta A, Jenkins AL, Lipinski MJ, Kim J, Chhour P, de Vries PS, Jinnouchi H, Kutys R, Mori H, Kutyna MD, Torii S, Sakamoto A, Choi CU, Cheng Q, Grove ML, Sawan MA, Zhang Y, Cao Y, Kolodgie FD, Cormode DP, Arking DE, Boerwinkle E, Morrison AC, Erdmann J, Sotoodehnia N, Virmani R, Finn AV. Cd163+ macrophages promote angiogenesis and vascular permeability accompanied by inflammation in atherosclerosis. J Clin Invest. 2018;128:1106–1124

37. Kristiansen M, Graversen JH, Jacobsen C, Sonne O, Hoffman HJ, Law SK, Moestrup SK. Identification of the haemoglobin scavenger receptor. Nature. 2001;409:198–201

38. Burdo TH, Lo J, Abbara S, Wei J, DeLelys ME, Preffer F, Rosenberg ES, Williams KC, Grinspoon S. Soluble cd163, a novel marker of activated macrophages, is elevated and associated with noncalcified coronary plaque in hiv-infected patients. J Infect Dis. 2011;204:1227–1236

39. Sánchez-Torres C, Gómez-Puertas P, Gómez-del-Moral M, Alonso F, Escribano JM, Ezquerra A, Domínguez J. Expression of porcine cd163 on monocytes/macrophages correlates with permissiveness to african swine fever infection. Arch Virol. 2003;148:2307–2323

40. Calvert JG, Slade DE, Shields SL, Jolie R, Mannan RM, Ankenbauer RG, Welch SK. Cd163 expression confers susceptibility to porcine reproductive and respiratory syndrome viruses. J Virol. 2007;81:7371–7379

41. Maidji E, Somsouk M, Rivera JM, Hunt PW, Stoddart CA. Replication of cmv in the gut of hiv-infected individuals and epithelial barrier dysfunction. PLoS Pathog. 2017;13:e1006202

42. Vita S, Lichtner M, Marchetti G, Mascia C, Merlini E, Cicconi P, Vullo V, Viale P, Costantini A, D’Arminio Monforte A, Group fIFS. Brief report: Soluble cd163 in cmv-infected and cmv-uninfected subjects on virologically suppressive antiretroviral therapy in the icona cohort. J Acquir Immune Defic Syndr. 2017;74:347–352

43. Foussat A, Bouchet-Delbos L, Berrebi D, Durand-Gasselin I, Coulomb-L’Hermine A, Krzysiek R, Galanaud P, Levy Y, Emilie D. Deregulation of the expression of the fractalkine/fractalkine receptor complex in hiv-1-infected patients. Blood. 2001;98:1678–1686

44. Sacre K, Hunt PW, Hsue PY, Maidji E, Martin JN, Deeks SG, Autran B, McCune JM. A role for cytomegalovirus-specific cd4+cx3cr1+t cells and cytomegalovirus-induced t-cell immunopathology in hiv-associated atherosclerosis. AIDS. 2012;26:805–814

45. Lee M, Lee Y, Song J, Lee J, Chang SY. Tissue-specific role of cx. Immune Netw. 2018;18:e5

46. McDermott DH, Halcox JP, Schenke WH, Waclawiw MA, Merrell MN, Epstein N, Quyyumi AA, Murphy PM. Association between polymorphism in the chemokine receptor cx3cr1 and coronary vascular endothelial dysfunction and atherosclerosis. Circ Res. 2001;89:401–407

47. Moatti D, Faure S, Fumeron F, Amara Ml-W, Seknadji P, McDermott DH, Debré P, Aumont MC, Murphy PM, de Prost D, Combadière C. Polymorphism in the fractalkine receptor cx3cr1 as a genetic risk factor for coronary artery disease. Blood. 2001;97:1925–1928

48. Stolla M, Pelisek J, von Brühl ML, Schäfer A, Barocke V, Heider P, Lorenz M, Tirniceriu A, Steinhart A, Bauersachs J, Bray PF, Massberg S, Schulz C. Fractalkine is expressed in early and advanced atherosclerotic lesions and supports monocyte recruitment via cx3cr1. PLoS One. 2012;7:e43572

49. Combadière B, Faure S, Autran B, Debré P, Combadière C. The chemokine receptor cx3cr1 controls homing and anti-viral potencies of cd8 effector-memory t lymphocytes in hiv-infected patients. AIDS. 2003;17:1279–1290

50. Hoehl S, Berger A, Ciesek S, Rabenau HF. Thirty years of cmv seroprevalence-a longitudinal analysis in a german university hospital. Eur J Clin Microbiol Infect Dis. 2020;39:1095–1102

51. Ahn J, Barber GN. Sting signaling and host defense against microbial infection. Exp Mol Med. 2019;51:1–10

52. Zhong B, Yang Y, Li S, Wang YY, Li Y, Diao F, Lei C, He X, Zhang L, Tien P, Shu HB. The adaptor protein mita links virus-sensing receptors to irf3 transcription factor activation. Immunity. 2008;29:538–550

53. Ishikawa H, Barber GN. Sting is an endoplasmic reticulum adaptor that facilitates innate immune signalling. Nature. 2008;455:674–678

54. Yamamoto T, Kanuma T, Takahama S, Okamura T, Moriishi E, Ishii KJ, Terahara K, Yasutomi Y. Sting agonists activate latently infected cells and enhance siv-specific responses ex vivo in naturally siv controlled cynomolgus macaques. Sci Rep. 2019;9:5917

55. Casazza JP, Brenchley JM, Hill BJ, Ayana R, Ambrozak D, Roederer M, Douek DC, Betts MR, Koup RA. Autocrine production of beta-chemokines protects cmv-specific cd4 t cells from hiv infection. PLoS Pathog. 2009;5:e1000646

