## Supplemental Tables for "Digital spatial profiling of coronary plaques from persons living with HIV reveals high levels of STING and CD163 in macrophage enriched regions"

Supplementary Tables: 2

**Supplementary Table 1. Demographics of HIV-negative and HIV-positive persons**

| | HIV-Positive n=6 | HIV-Negative n=6 |
|---|---|---|
| **Age, yrs.** | 50 [47, 66] | 52 [48, 65] |
| **Gender, % Male** | 3 | 3 |
| **Race, % Caucasian** | 1 | 5 |
| | **Plaque characteristics** | |
| **Early atheroma, %** | 3 | 4 |
| **Late atheroma, %** | 3 | None |
| **Chronic Total Occlusion, %** | None | 1 (17%) |
| **Non-specified atheroma, %** | None | 1 (17%) |
| **Vessel type, number** | PLAD (2)<br>PRC (1)<br>PRC1 (1)<br>LOM1 (1)<br>MLAD (MRC) (1) | PRC1 (2)<br>PRC2 (1)<br>MRC3 (1)<br>PLC11 (1)<br>PLAD (1) |
| **Sudden death category** | Non-cardiac death (1)<br>Cardiac death (but non-CAD) (4)<br>Stent/CABG (1) | Severe CAD w/o acute thrombus (3)<br>Erosion (1)<br>Rupture (1)<br>Non-cardiac death (1) |

*Abbreviation: yrs, years; PLAD, proximal left anterior descending; PRC, proximal right coronary artery; LOM, left obtuse marginal; MLAD, middle left anterior descending coronary artery; MRC, MRC middle right coronary; CABG, coronary artery bypass, non-CAD, non-coronary artery disease*

**Supplementary Table 2. Demographics of HIV-negative and HIV-positive persons used in digital spatial profiling**

| | HIV-positive (n=1) | HIV-Negative (n=1) |
|---|---|---|
| **Age, yrs.** | 49 | 51 |
| **Gender** | Male | Male |
| **Race** | African American | Hispanic |
| **Vessel** | PRC | PLC11 |
| **Plaque Type** | Early Fibroatheroma | Chronic total occlusion |
| **Plaque Area, μm²** | $7.65E^6$ | $2.82E^6$ |
| **% stenosis** | 59.95 | 73.72 |
| **Immunohistochemistry stains** | | |
| **% CD68** | 2.70 | 2.33 |
| **% CD163** | 1.45 | 0.35 |
| **% CD3** | 0.52 | 0.42 |
| **% CD4** | 0.16 | 0.07 |
| **% CD8** | 0.35 | 0.15 |
| **% VCAM-1** | 0.23 | 0.07 |
| **% CX3CR1** | 1.37 | 1.99 |

**Abbreviations: yrs., years; PRC, proximal right coronary artery; PLC, posterior circumflex branch (obtuse marginal); vascular cell adhesion molecule, VCAM-1**
