## Supplemental Figures for "Digital spatial profiling of coronary plaques from persons living with HIV reveals high levels of STING and CD163 in macrophage enriched regions"

Supplementary Figures: 2

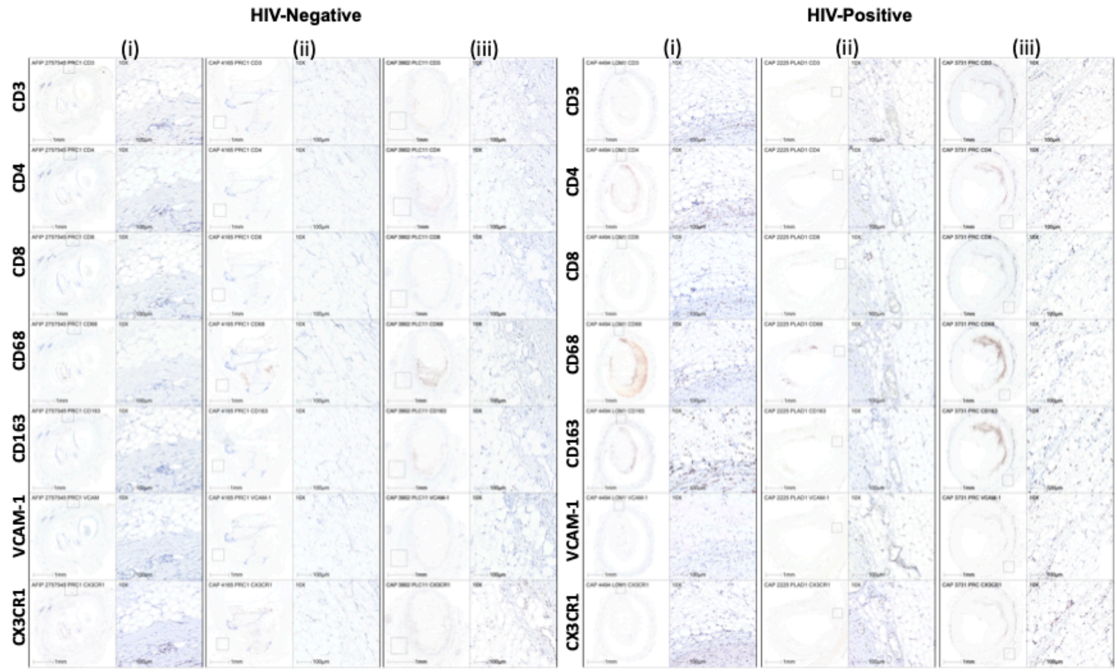


**Supplementary Figure 1. Innate and adaptive immune cells within the perivascular adipose tissue present in coronary plaques of PLWH.** Representative images of immunohistochemical stains from HIV-positive (n=3) and HIV-negative (n=3) perivascular adipose tissue adjacent to coronary plaques.

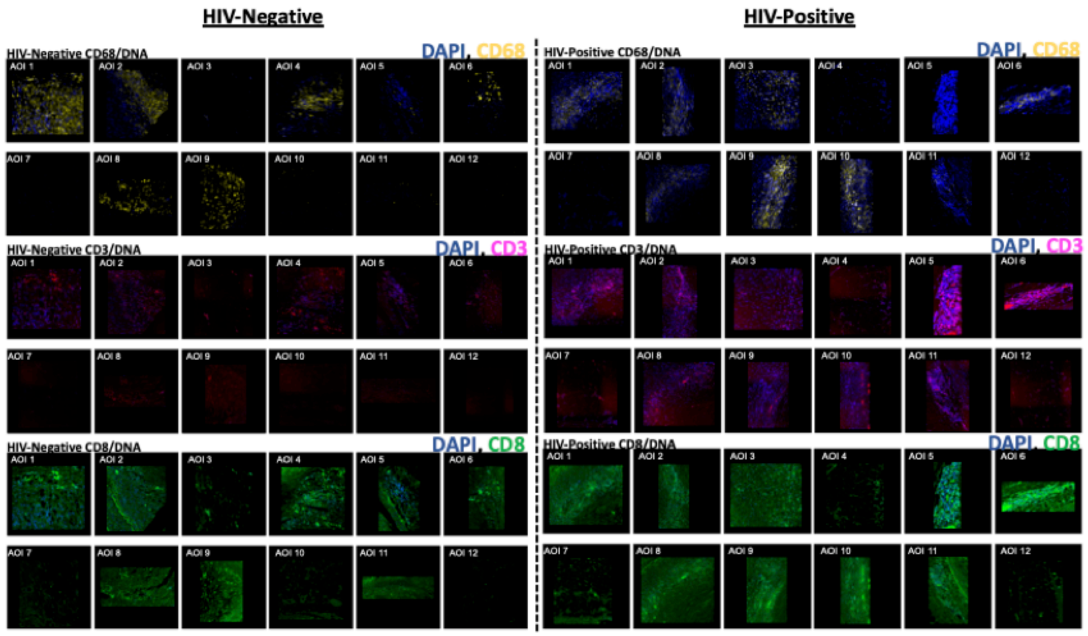


**Supplementary Figure 2. Fluorescent imaging showing different AOIs within the coronary plaque.** Dual-color fluorescent images of AOIs within the coronary plaque are shown in the figure. Six-representative segments out of a total of 7 are shown in the HIV-negative sample and 6/6 in the HIV-positive sample.
